# IKBIP is a novel EMT-related biomarker and predicts poor survival in glioma

**DOI:** 10.1101/2020.06.26.172569

**Authors:** Ying Yang, Jin Wang, Shihai Xu, Wen Lv, Fei Shi, Aijun Shan

## Abstract

**Purpose:** I kappa B-interacting protein (IKBIP) in cancer has rarely been reported. This study aimed at investigating its expression pattern and biological function in brain glioma at transcriptional level.

**Methods:** We selected 301 glioma patients with microarray data from CGGA database and 697 glioma patients with RNAseq data from TCGA database. Transcriptome data and clinical data of 998 samples were analyzed. Statistical analysis and figure generating were performed with R language.

**Results:** We found that IKBIP expression showed positive correlation with WHO grade of glioma. IKBIP was increased in IDH wildtype and mesenchymal molecular subtype of glioma. Gene ontology analysis demonstrated that IKBIP was profoundly associated with extracellular matrix organization, cell-substrate adhesion and response to wounding in both pan-glioma and glioblastoma. Subsequent GSEA analysis revealed that IKBIP was particularly correlated with epithelial-to-mesenchymal transition (EMT). To further elucidate the relationship between IKBIP and EMT, we performed GSVA analysis to screen the EMT-related signaling pathways, and found that IKBIP expression was significantly associated with PI3K/AKT, hypoxia and TGF-β pathway. Moreover, IKBIP expression was found to be synergistic with key biomarkers of EMT, especially with N-cadherin, vimentin, snail, slug and TWIST1. Finally, higher IKBIP indicated significantly shorter survival for glioma patients.

**Conclusions:** IKBIP was associated with more aggressive phenotypes of gliomas.

Furthermore, IKBIP was significantly involved in EMT and could serve as an independent prognosticator in glioma.

## 1. Introduction

Gliomas account for the most common and aggressive primary brain cancers among adult patients^[1]^. Despite great advances in diagnosis and treatment, the prognosis for glioma patients remains unfavorable. Especially for those with glioblastoma (GBM), the most devastating type, the median survival time is only about fifteen months^[2, 3]^. Epithelial-to-mesenchymal transition (EMT) has been widely reported as a key mechanism in promoting migration, invasion and tumor progression in glioma^[4]^. Identification of novel EMT-related markers is of great necessity.

I kappa B-interacting protein (IKBIP) was found to be one of the target genes of p53, and play a crucial role in proapoptotic function^[5]^. Recently, IKBIP was identified as a vital modulator of inflammation^[6]^. Heretofore, the biological function of IKBIP in malignancies has been rarely reported. Only one study^[7]^, through weighted gene co-expression network analysis (WGCNA), preliminarily revealed IKBIP as a potential hub gene in gliomagenesis. However, the role of IKBIP in glioma still remains largely unclear. In the present study, we took advantage of 998 glioma patients with transcriptome data to investigate the clinical significance, molecular characterization and biological function of IKBIP in glioma.

## 2. Materials and Methods

### 2.1. Sample and data collection

Transcriptome and clinical data of glioma patients were available in Chinese Glioma Genome Atlas (CGGA) website (http://www.cgga.org.cn/) and TCGA website (http://cancergenome.nih.gov/). In total, 998 glioma patients, including 301 CGGA microarray data (GeneSpring GX 11.0 normalization) and 697 TCGA RNAseq data (RSEM normalization, level 3), were enrolled. The baseline characteristics of patients in both cohorts were described in Table S1.

### 2.2. Statistical analysis

For TCGA cohort, RSEM RNAseq data were log2 transformed. For CGGA cohort, microarray data (already normalized and centered by data provider) were directly analyzed. Statistical analysis was performed with R language. Multiple R packages, including ggplot2, pROC^[8]^, pheatmap, corrgram, circlize^[9]^, gsva, as well as survival, were used to generate figures. The biological processes of IKBIP-related genes were annotated using Metascape^[10]^ (https://metascape.org). Hallmark gene sets were downloaded from Gene Set Enrichment Analysis (GSEA) website (http://software.broadinstitute.org/) for GSEA^[11]^ and Gene Set Variation Analysis (GSVA)^[12]^. All statistical tests were two-sided, and a *p* value of < 0.05 was considered as a statistical significance.

## 3. Results

### 3.1. IKBIP expression was correlated with aggressive phenotypes of glioma

IKBIP expression levels were compared across different WHO grades. The results of both CGGA and TCGA cohorts consistently showed a significant positive correlation between IKBIP expression and WHO grade, except for comparison between WHO II and WHO III in CGGA, which also showed apparent trend (Figs. 1A and E). In addition, when patients were subclassified with respect to IDH mutation status, IDH wildtype was found to be more associated with an increased pattern of IKBIP expression in both datasets, even though no statistical significance was reached in some groups (Figs. 1B and F). These results suggested that higher IKBIP was paralleled with higher malignancy in glioma. Moreover, the correlation between IKBIP and TCGA molecular subtype was further examined. As shown in Figs. 1C and G, IKBIP expression in mesenchymal subtype was significantly upregulated than that in other subtypes, suggesting that IKBIP expression could contribute as a specific marker for mesenchymal subtype. ROC curves were subsequently performed to evaluate the performance of IKBIP in distinguishing mesenchymal subtype. Areas under curves (AUC) were 88.5% in CGGA and 85.5% in TCGA, respectively (Figs. 1D and H).

**Fig. 1.**
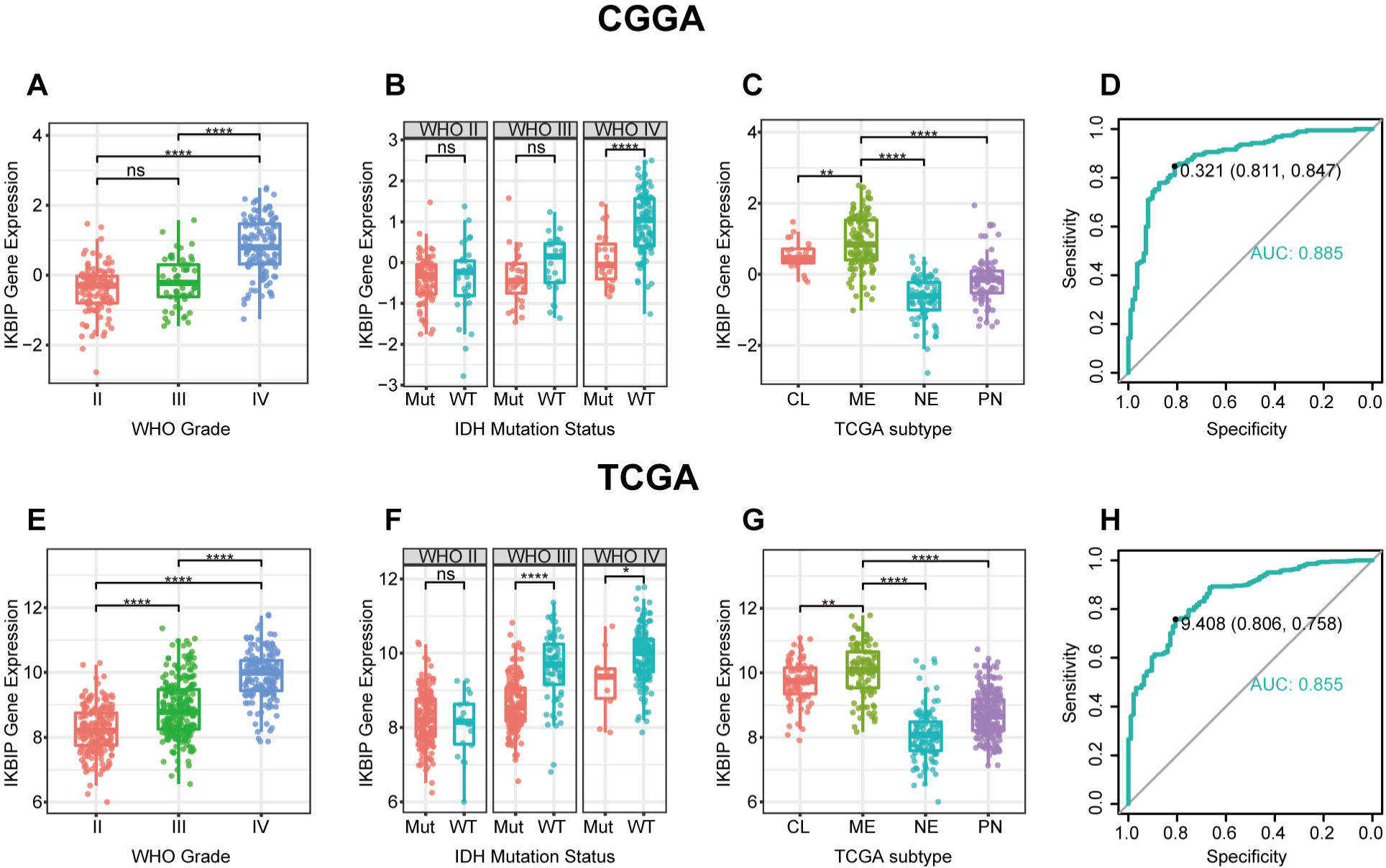
IKBIP expression in CGGA and TCGA dataset according to WHO grade (A, E), IDH mutation status (B, F), TCGA molecular subtype (C, G) and ROC curves (D, H) for distinguishing mesenchymal subtype. * indicates p value < 0.05, **indicates p value < 0.01, *** indicates p value < 0.001, **** indicates p value < 0.0001.

### 3.2. IKBIP-related biological process

To explore the biological process of IKBIP in glioma, Pearson correlation test was performed between IKBIP and other genes. With the criteria of Pearson coefficient |r| > 0.6, we identified 711 IKBIP-positively-correlated genes and 462 IKBIP-negatively-correlated genes in CGGA, and 938 IKBIP-positively-correlated genes and 17 IKBIP-negatively-correlated genes in TCGA. To ensure accuracy, IKBIP-significantly-correlated genes that were overlapped between both datasets were selected for GO analysis. A venn diagram (Fig. S1A) was constructed, illustrating an overlap of 376 IKBIP-positively-correlated genes (Table. S2), which were subsequently annotated with GO analysis. We found that IKBIP-positively-correlated genes were mainly involved in EMT-related biological processes including extracellular structure organization, cell-substrate adhesion, blood vessel development, response to wounding, and response to growth factor. Other biological processes included neutrophil degranulation and cell division, pointing toward an association between IKBIP and regulation of immune response and cell cycle, respectively (Figs. 2A and B).

**Fig. 2.**
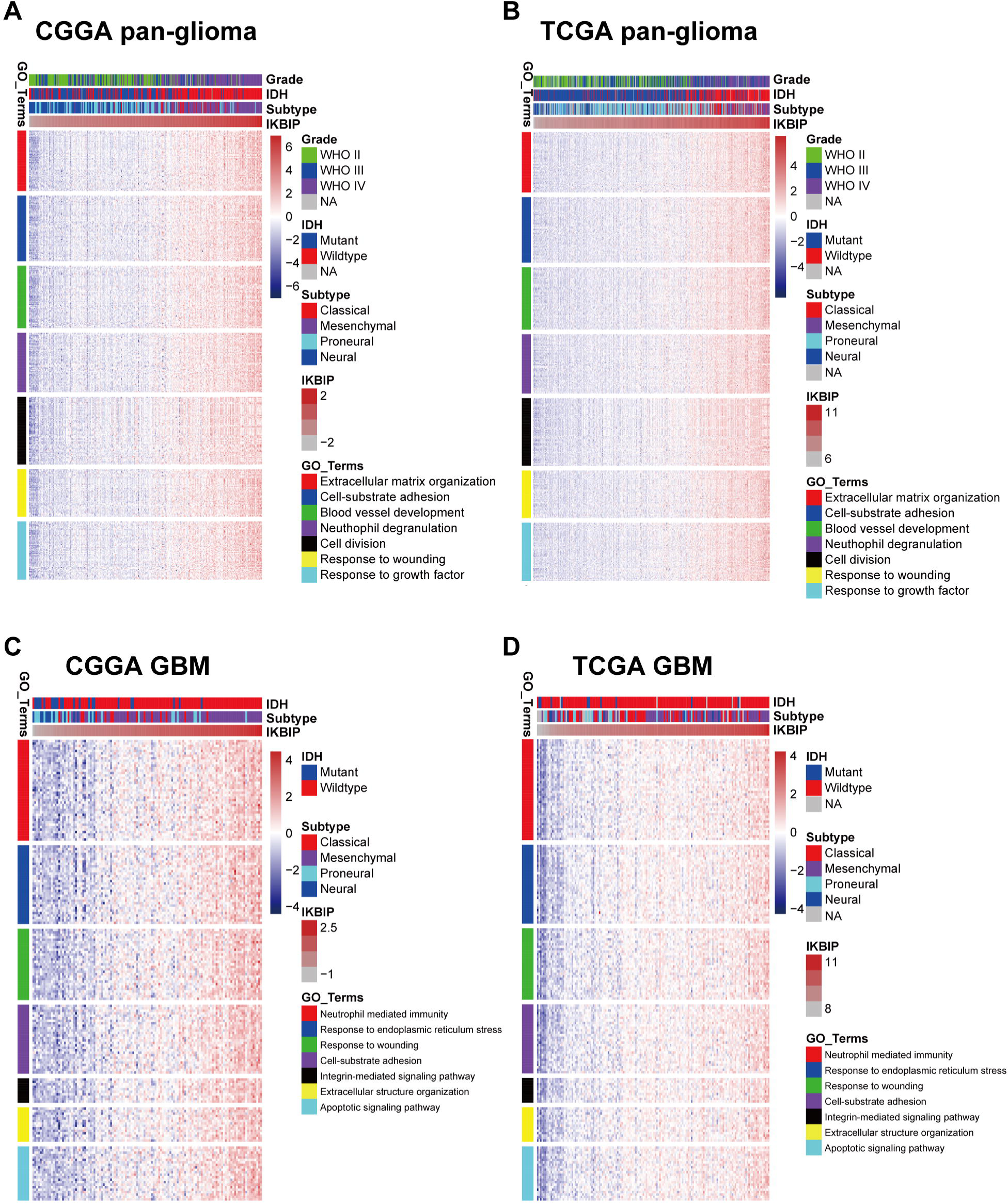
Gene Ontology analysis for IKBIP in pan-glioma (A, B) and GBM (C, D).

In view of GBM as a distinct subgroup of glioma, we then conducted an independent GO enrichment analysis in this group. In GBM of both datasets, venn diagram (Fig. S1B) exhibited an overlap of 191 IKBIP-positively-correlated genes (Table. S3). Subsequent GO analysis revealed that these genes were also significantly associated with EMT-related biological processes including response to endoplasmic reticulum stress^[13]^, response to wounding, cell-substrate adhesion, integrin-mediated signaling pathway^[14]^, and extracellular structure organization. Besides, IKBIP seemed to be more associated with neutrophil mediated immunity, suggesting that IKBIP upregulation was accompanied by immunosuppression of GBM, which indicated a more malignant characteristic in glioma. While it should be noted that IKBIP showed positive correlation with apoptotic signaling pathway, enlightening us that IKBIP might also act as a pro-apoptotic factor in GBM^[5]^ (Figs. 2C and D).

### 3.3. IKBIP was associated with EMT

GSEA analyses were performed in both CGGA and TCGA datasets, and it turned out that, IKBIP was significantly correlated with EMT in CGGA (NES = 1.968, FDR = 0.010) (Fig. 3A), which was further validated in TCGA (NES = 1.747, FDR = 0.058) (Fig. 3B). Furthermore, in GBM, IKBIP showed an even higher association with EMT in both cohorts (Figs. 3C and D). These results indicated that IKBIP could be profoundly associated with EMT phenotype in glioma.

**Fig. 3.**
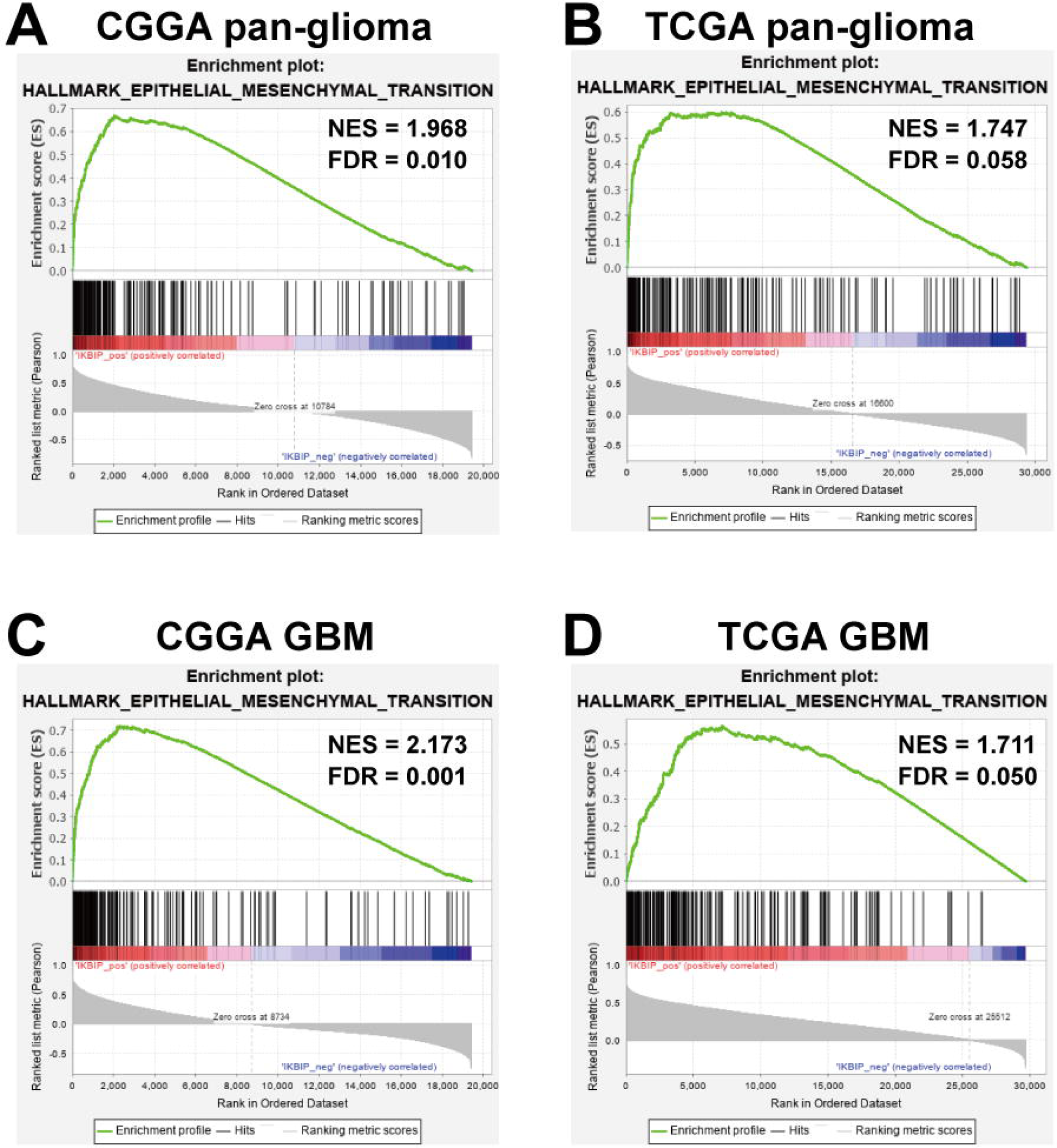
Gene set enrichment analysis (GSEA) for enrichment of EMT according to IKBIP expression in pan-glioma (A, B) and GBM (C, D).

### 3.4. IKBIP interacted with EMT-related signaling pathways in glioma

To further investigate the relationship between IKBIP and EMT, we downloaded seven gene sets from GSEA website (Table S4), which were subsequently transformed into metagenes, representing different EMT-related signaling pathways, summarized by Gonzalez et al^[15]^. As shown in Figs. 4A and B, three clusters including TGF-β, PI3K/AKT, and hypoxia signaling pathway, were significantly associated with IKBIP expression. To quantify what we observed in clusters, Gene Set Variation Analysis (GSVA) was performed to generate seven metagenes based on corresponding genes of seven EMT-related signaling pathways. According to Pearson r value between IKBIP and seven metagenes, Corrgrams were generated to evaluate their intercorrelations (Figs. 4C and D). IKBIP showed a robust correlation with TGF-β, PI3K/AKT, and hypoxia signaling pathway, while only showed a weak correlation with WNT, MAPK, NOTCH, and HEDGEHOG pathway, in consistent with what we observed in Figs 4A and B. Moreover, a similar pattern of EMT-related signaling pathways was also observed in GBM of both CGGA and TCGA dataset (Fig. 5).

**Fig. 4.**
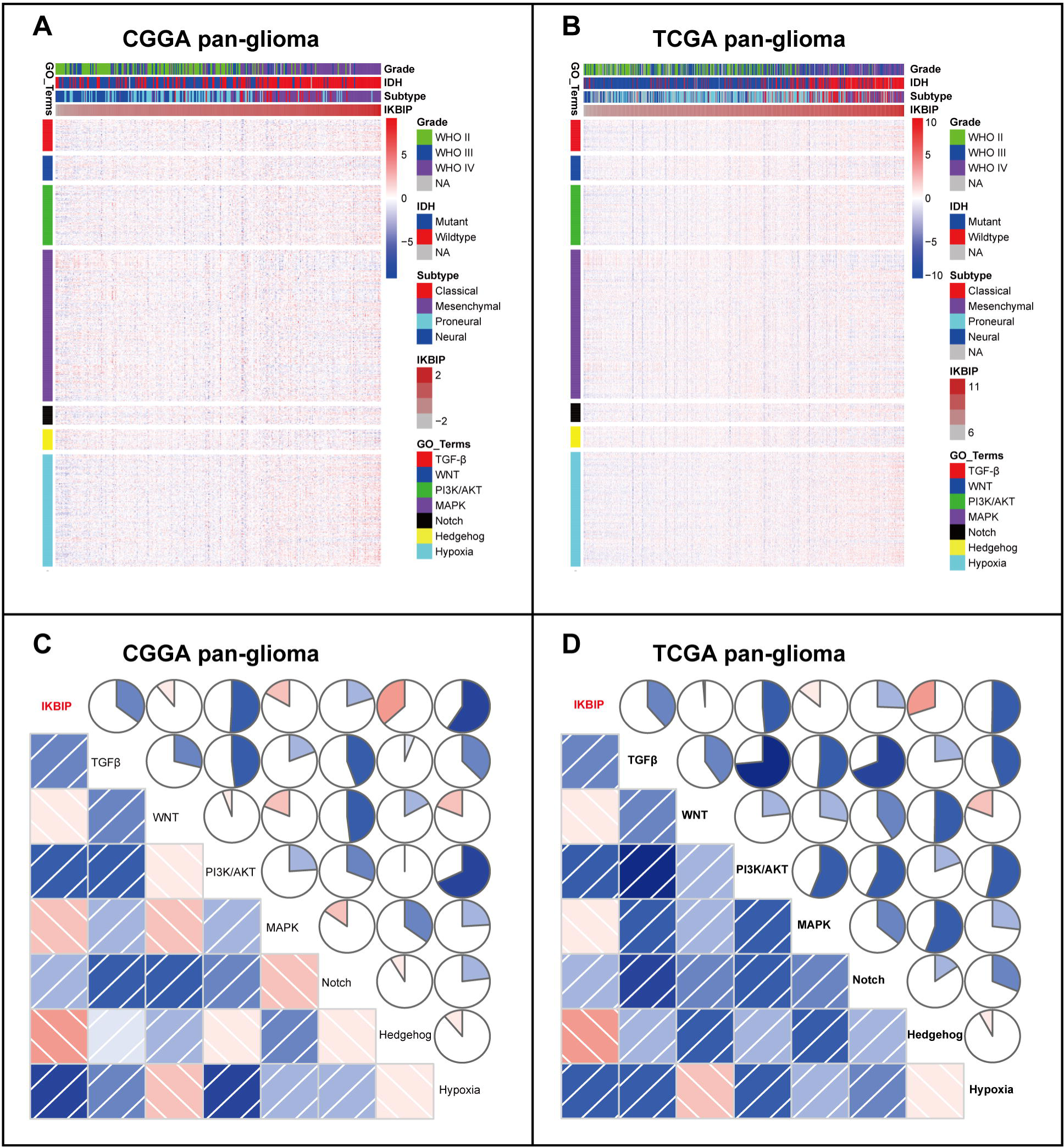
Cluster (A, B) and GSVA (C, D) of IKBIP-related EMT signaling pathways in pan-glioma.

**Fig. 5.**
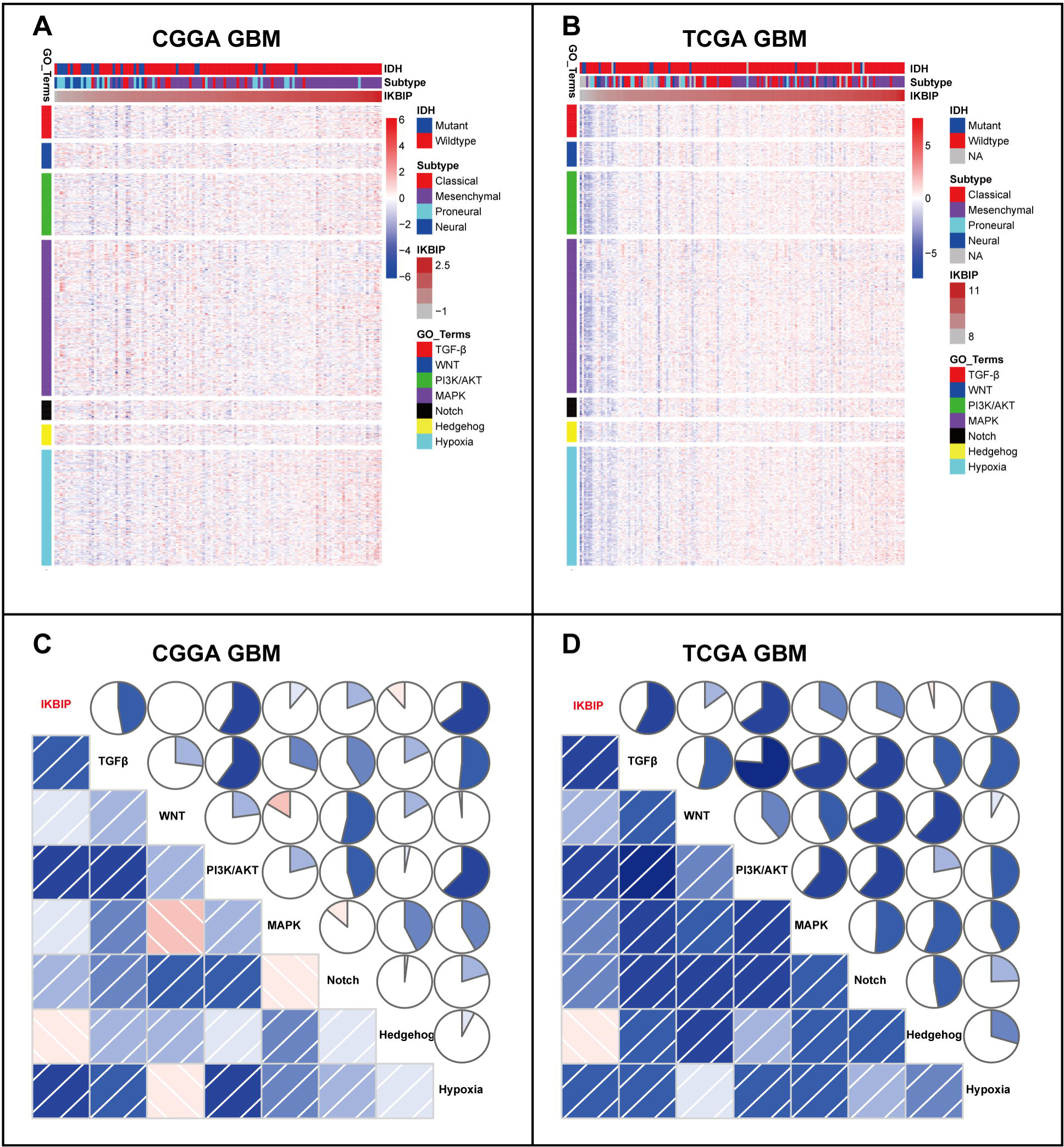
Cluster (A, B) and GSVA (C, D) of IKBIP-related EMT signaling pathways in GBM.

### 3.5. IKBIP interacted with EMT-related key biomarkers in glioma

To further validate the role of IKBIP in EMT-related signaling pathways, we examined the correlation between IKBIP and EMT-related key biomarkers including E-cadherin, N-cadherin, vimentin, snail and slug. Circos plots revealed that IKBIP expression was significantly associated with N-cadherin, vimentin, snail and slug (Figs. 6A and B). To further demonstrate the interaction of these markers in GBM, Pearson correlation tests were additionally performed. As shown in Figs. 6C and D, the correlation between IKBIP and these markers in GBM was also very robust in both datasets, indicating synergistic effects of these members during glioma EMT. While the correlation between IKBIP and E-cadherin was very weak, which might be deemed as a noise.

**Fig. 6.**
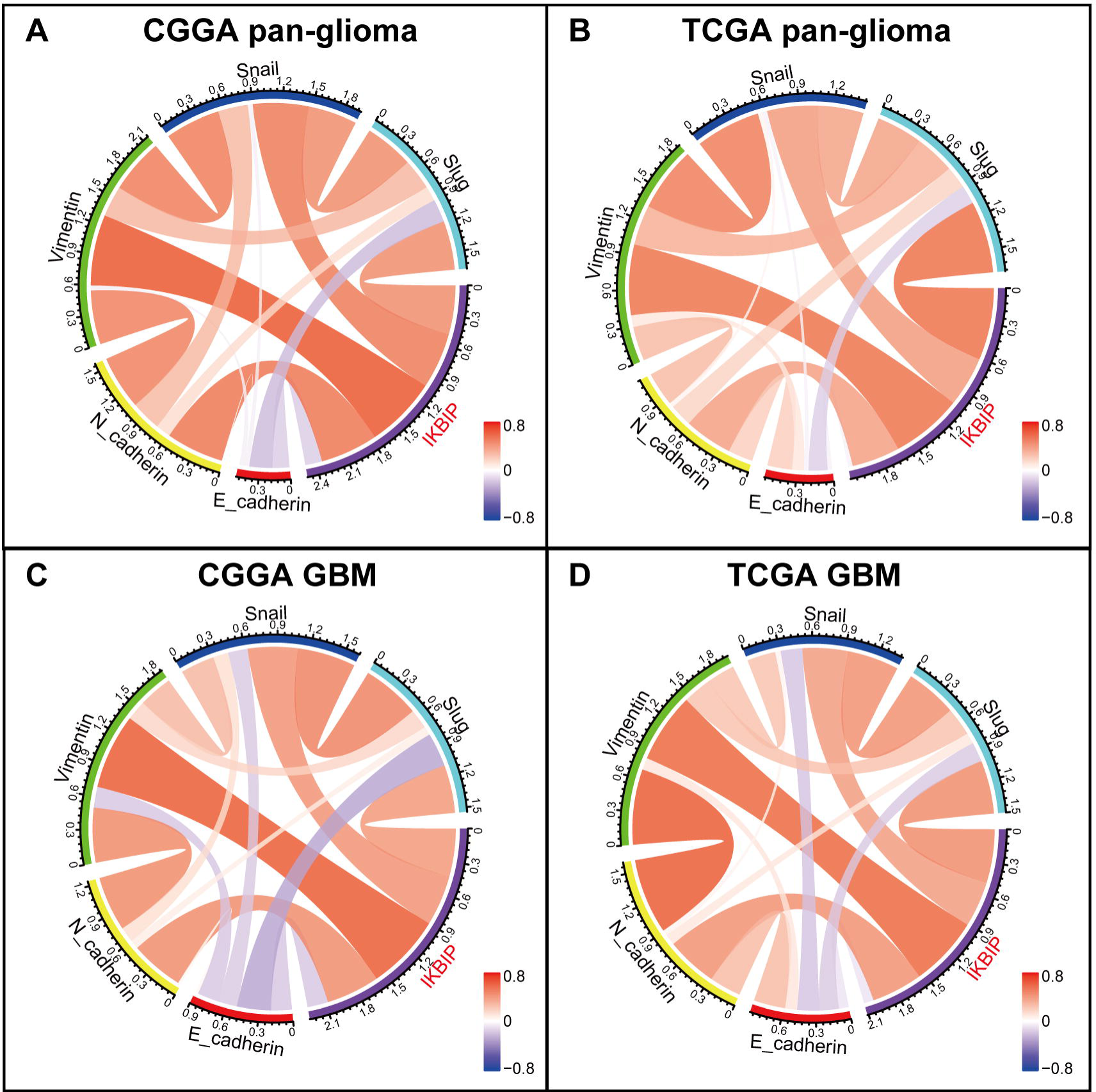
Correlation of IKBIP and EMT key biomarkers.

There are many other molecules that have been identified as EMT-related key biomarkers in EMT^[16]^. We additionally enrolled EMT-related markers including ZEB1/2, β-catenin, and TWIST1/2, and put them into analysis together with IKBIP. Subsequent Circos plots in both CGGA and TCGA congruently revealed that IKBIP expression was especially correlated with TWIST1 in both pan-glioma and GBM (Fig. 7).

**Fig. 7.**
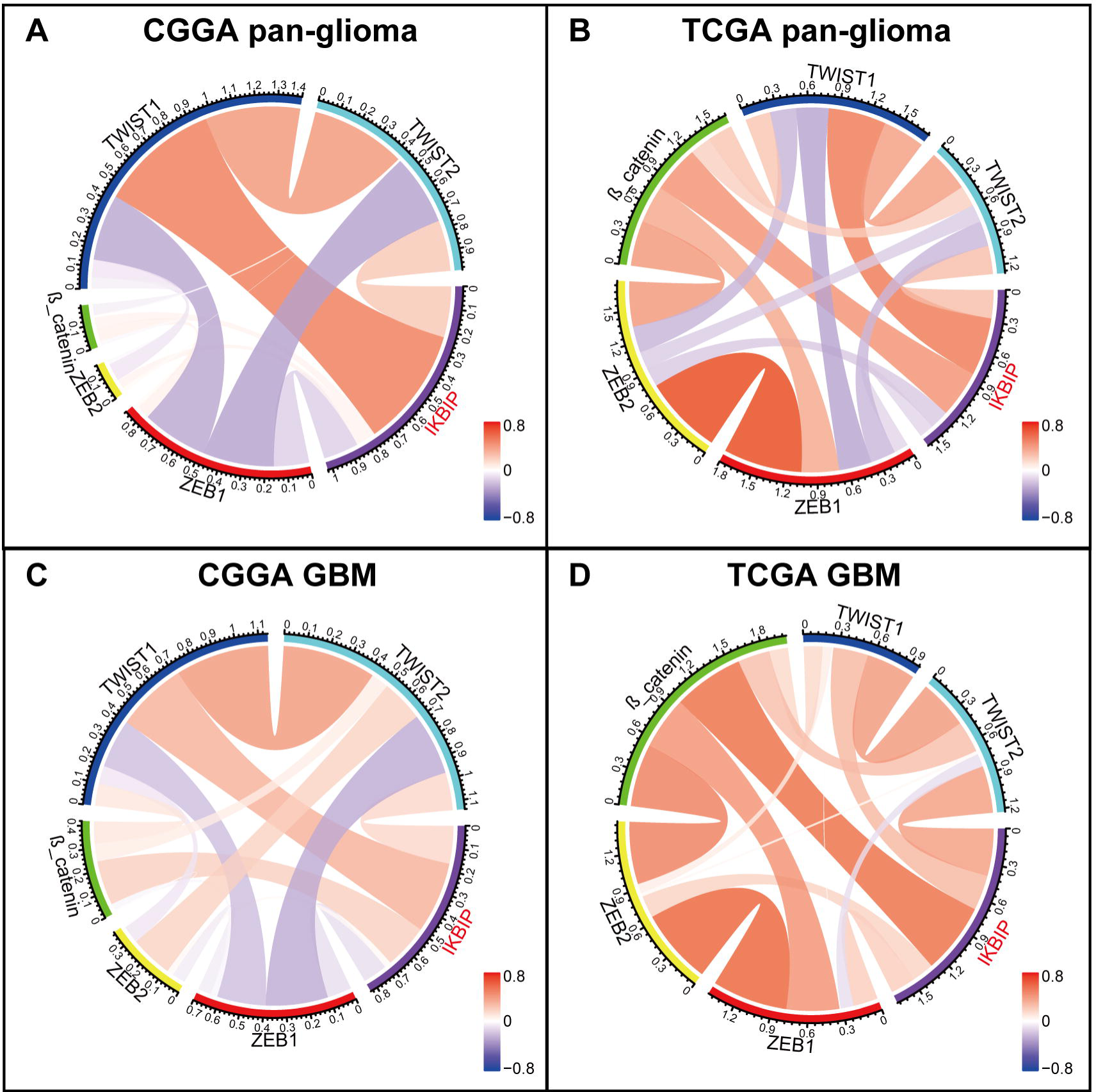
Correlation of IKBIP and other EMT key biomarkers.

### 3.6. Higher IKBIP predicts shorter survival for glioma

Kaplan-Meier (KM) survival analyses were performed to examine the prognostic value of IKBIP in glioma. According to IKBIP expression, pan-glioma samples were divided into two groups in each dataset. As shown in Figs. 8A and D, a higher level of IKBIP expression predicted a significantly shorter survival. Moreover, a similar pattern of the KM survival curve was observed among patients with LGG (Figs. 8B and E) and GBM (Figs. 8C and F).

**Fig. 8.**
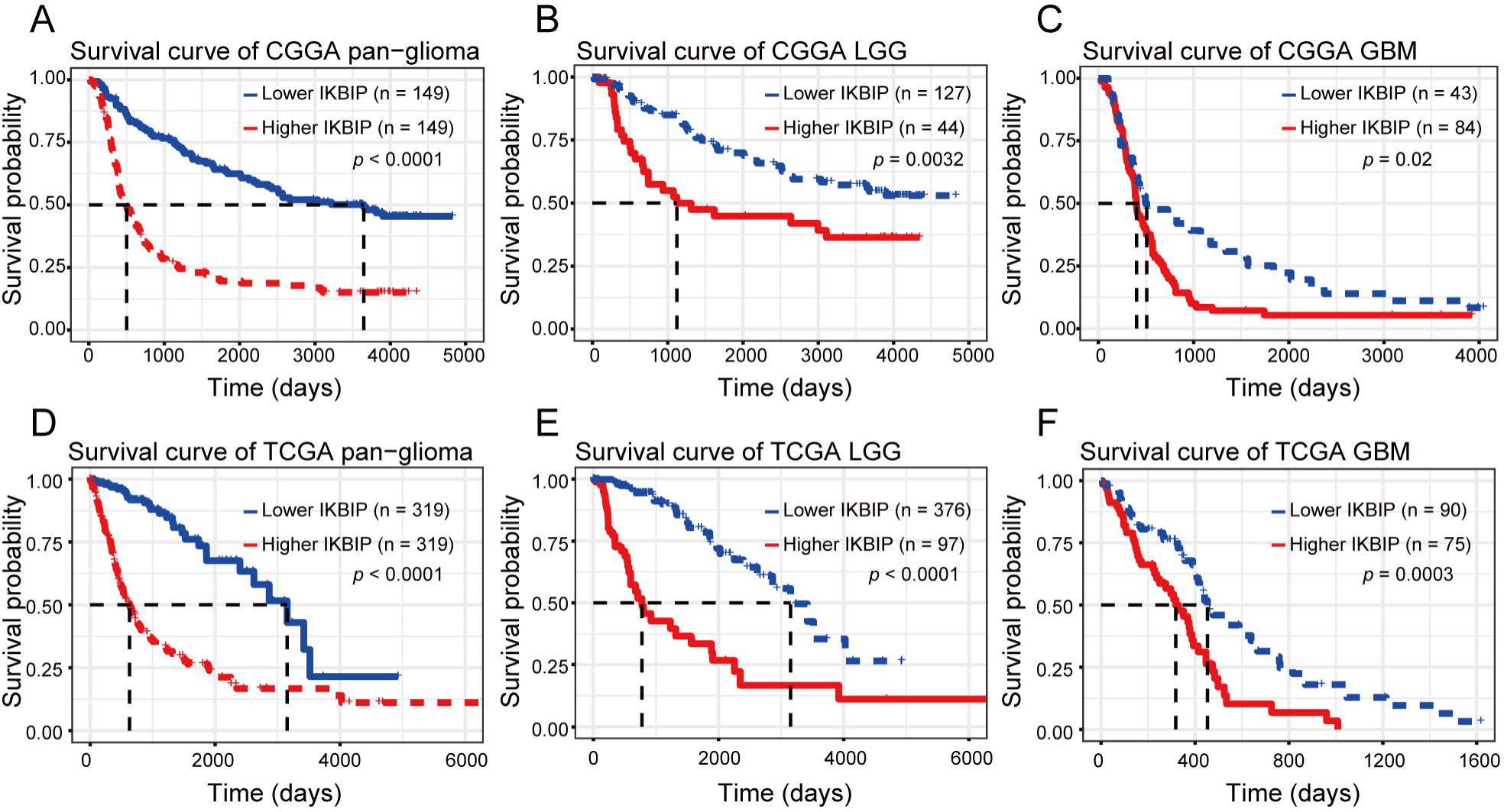
Survival analysis for IKBIP in pan-glioma (A, D), LGG (B, E) and GBM (C, F).

## 4. Discussion

In the present study, we investigated the transcriptional expression profiles of IKBIP in 998 glioma patients and revealed that IKBIP expression showed significantly positive correlation with WHO grade of glioma. Furthermore, higher IKBIP expression was usually accompanied by a more aggressive and malignant phenotype in glioma, including GBM, IDH wildtype and mesenchymal subtype. Moreover, higher IKBIP expression indicated a significantly shorter survival for patients with glioma, across different WHO grades. These findings suggested that IKBIP played a vital role in the malignant progression of gliomas, in line with what Chen et al. found in their WGCNA study^[7]^. Understanding the molecular mechanism of IKBIP in glioma may provide a novel therapeutic target to overcome this fatal disease.

To elucidate the biological function of IKBIP in glioma, GO analysis was performed, and it turned out that, IKBIP was highly associated with a series of EMT-related biological processes including extracellular matrix organization, cell-substrate adhesion, and response to wounding in both pan-glioma and GBM. Subsequent GSEA analysis revealed remarkable evidence for that IKBIP was particularly correlated with EMT, which had been extensively confirmed to play a key role not only in glioma migration/invasion but also in glioma recurrence and therapeutic resistance^[17-19]^. These results enlightened us that IKBIP might promote tumorigenesis and progression of glioma mainly by means of EMT induction, which has yet been previously reported. Besides, GO analysis also revealed that IKBIP played a crucial role in tumor-induced immune and inflammatory response in glioma, especially in GBM, in line with the results presented by Wu et al^[6]^. They demonstrated that IKBIP played an inhibitory role in immune and inflammatory response through negative regulation of NF-kappaB pathway. Based on these, we concluded that, other than as a key molecule for EMT induction, IKBIP might contribute as an immune suppressor in glioma as well, which further validated its oncogenic role in glioma. Meanwhile, it was noteworthy that IKBIP showed robust correlation with apoptotic signaling pathway in GBM, suggesting a potential proapoptotic function^[5]^. As a result, we speculated that IKBIP might have a dualistic nature in gliomagenesis, and the robust pro-tumoral effect through EMT induction and immune inhibition overwhelmed the anti-tumoral effect through proapoptotic function.

To further validate the pro-EMT effect of IKBIP in glioma, we selected a series of EMT-related signaling pathways and biomarkers, which were then analyzed to determine their interaction with IKBIP, and found that IKBIP showed robust correlation with PI3K/AKT, hypoxia, and TGF-β signaling pathway, suggesting that IKBIP might promote EMT process through these pathways. Moreover, most of EMT biomarkers including N-cadherin, snail, slug, vimentin, and TWIST1 were significantly associated with IKBIP, indicating that IKBIP interacted synergistically with these key molecules of EMT. These results further validated the involvement of IKBIP in glioma EMT. Thus, our findings might bring a novel EMT target for potential glioma treatment.

## Conclusions

In conclusion, IKBIP expression was associated with more aggressive phenotypes of glioma and predicted much worse survival for patients. Moreover, IKBIP was significantly associated with EMT process, and interacted synergistically with EMT-related signaling pathways and key biomarkers. However, a limitation of the current study was that no experimental validation was performed. Further *in-vitro* and *in-vivo* studies are needed to validate its role in glioma.

## Acknowledgments

We appreciate the generosity of CGGA project and TCGA project for sharing data. This work was supported by Medical Scientific Research Foundation of Shenzhen Health Commission (szfz2018022) and Science and Technology Innovation Foundation of Shenzhen (JCYJ20190806150005453).

## Figure Legends

**Fig. S1** Venn Diagram of IKBIP-positively-correlated genes in pan-glioma (A) and GBM (B).

## References

[1] Meng X, Zhao Y, Han B, Zha C, Zhang Y, Li Z, Wu P, et al. Dual functionalized brain-targeting nanoinhibitors restrain temozolomide-resistant glioma via attenuating EGFR and MET signaling pathways. Nat Commun. 2020; 11:594.

[2] Yang W, Wu PF, Ma JX, Liao MJ, Wang XH, Xu LS, Xu MH, et al. Sortilin promotes glioblastoma invasion and mesenchymal transition through GSK-3β/β-catenin/twist pathway. Cell Death Dis. 2019; 10:208.

[3] Wei J, Ouyang X, Tang Y, Li H, Wang B, Ye Y, Jin M, et al. ER-stressed MSC displayed more effective immunomodulation in RA CD4(+)CXCR5(+)ICOS(+) follicular helper-like T cells through higher PGE2 binding with EP2/EP4. Mod Rheumatol. 2019; 30:509–16.

[4] Ma YS, Wu ZJ, Bai RZ, Dong H, Xie BX, Wu XH, Hang XS, et al. DRR1 promotes glioblastoma cell invasion and epithelial-mesenchymal transition via regulating AKT act ivation. Cancer Lett. 2018; 423:86–94.

[5] Hofer-Warbinek R, Schmid JA, Mayer H, Winsauer G, Orel L, Mueller B, Wiesner C, et al. A highly conserved proapoptotic gene, IKIP, located next to the APAF1 gene locus, is regulated by p53. Cell Death Differ. 2004; 11:1317–25.

[6] Wu H, Liu H, Zhao X, Zheng Y, Liu B, Zhang L, Gao C. IKIP Negatively Regulates NF-kappaB Activation and Inflammation through Inhibition of IKKalpha/beta Phosphorylation. J Immunol. 2020; 204:418–27.

[7] Chen TY, Liu Y, Chen L, Luo J, Zhang C, Shen XF. Identification of the Potential Biomarkers in Patients with Glioma: A Weighted Gene Co-Expression Network Analysis. Carcinogenesis. 2019.

[8] Robin X, Turck N, Hainard A, Tiberti N, Lisacek F, Sanchez JC, Muller M. pROC: an open-source package for R and S+ to analyze and compare ROC curves. BMC Bioinformatics. 2011; 12:77.

[9] Gu Z, Gu L, Eils R, Schlesner M, Brors B. circlize Implements and enhances circular visualization in R. Bioinformatics. 2014; 30:2811–2.

[10] Zhou Y, Zhou B, Pache L, Chang M, Khodabakhshi AH, Tanaseichuk O, Benner C, et al. Metascape provides a biologist-oriented resource for the analysis of systems-level datasets. Nat Commun. 2019; 10:1523.

[11] Subramanian A, Tamayo P, Mootha VK, Mukherjee S, Ebert BL, Gillette MA, Paulovich A, et al. Gene set enrichment analysis: a knowledge-based approach for interpreting genome-wide expression profiles. Proc Natl Acad Sci U S A. 2005; 102:15545–50.

[12] Hanzelmann S, Castelo R, Guinney J. GSVA: gene set variation analysis for microarray and RNA-seq data. BMC Bioinformatics. 2013; 14:7.

[13] Shin HS, Ryu ES, Oh ES, Kang DH. Endoplasmic reticulum stress as a novel target to ameliorate epithelial-to-mesenchymal transition and apoptosis of human peritoneal mesothelial cells. Lab Invest. 2015; 95:1157–73.

[14] Park H, Kim D, Kim D, Park J, Koh Y, Yoon SS. Truncation of MYH8 tail in AML: a Novel prognostic marker with increase cell migration and epithelial-mesenchymal transition utilizing RAF/MAPK pathway. Carcinogenesis. 2019.

[15] Gonzalez DM, Medici D. Signaling mechanisms of the epithelial-mesenchymal transition. Sci Signal. 2014; 7:re8.

[16] Xu J, Zhang Z, Qian M, Wang S, Qiu W, Chen Z, Sun Z, et al. Cullin-7 (CUL7) is overexpressed in glioma cells and promotes tumorigenesis via NF-κB activation. J Exp Clin Cancer Res. 2020; 39:59.

[17] Li C, Zheng H, Hou W, Bao H, Xiong J, Che W, Gu Y, et al. Long non-coding RNA linc00645 promotes TGF-β-induced epithelial-mesenchymal transition by regulating miR-205-3p-ZEB1 axis in glioma. Cell Death Dis. 2019; 10:717.

[18] Li H, Li J, Chen L, Qi S, Yu S, Weng Z, Hu Z, et al. HERC3-Mediated SMAD7 Ubiquitination Degradation Promotes Autophagy-Induced EMT and Chemoresistance in Glioblastoma. Clin Cancer Res. 2019; 25:3602–16.

[19] Li H, Li J, Zhang G, Da Q, Chen L, Yu S, Zhou Q, et al. HMGB1-Induced p62 Overexpression Promotes Snail-Mediated Epithelial-Mesenchymal Transition in Gliobla stoma Cells via the Degradation of GSK-3β. Theranostics. 2019; 9:1909–22.

