## Supplementary figures and images for "IKBIP is a novel EMT-related biomarker and predicts poor survival in glioma"

### Supplemental Figure 1

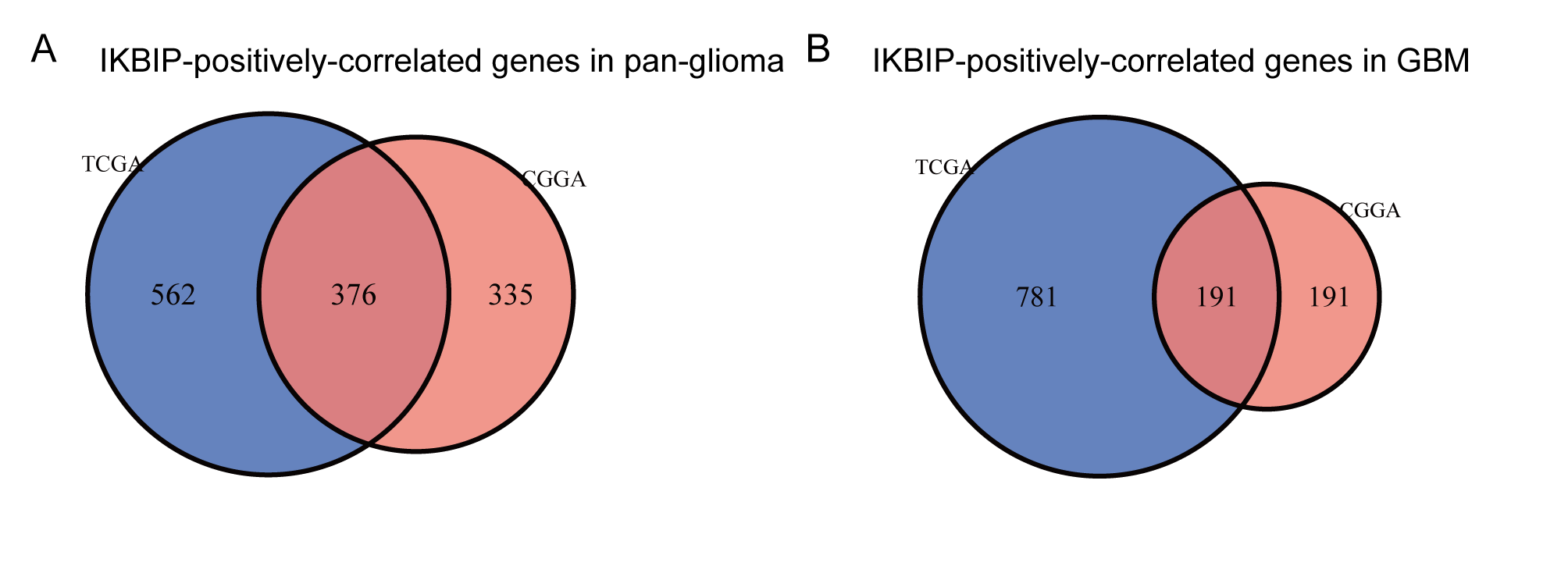
